# DHSpred: support-vector-machine-based human DNase I hypersensitive sites prediction using the optimal features selected by random forest

**DOI:** 10.1101/224527

**Authors:** Balachandran Manavalan, Tae Hwan Shin, Gwang Lee

## Abstract

DNase I hypersensitive sites (DHSs) are genomic regions that provide important information regarding the presence of transcriptional regulatory elements and the state of chromatin. Therefore, identifying DHSs in uncharacterized DNA sequences is crucial for understanding their biological functions and mechanisms. Although many experimental methods have been proposed to identify DHSs, they have proven to be expensive for genome-wide application. Therefore, it is necessary to develop computational methods for DHS prediction. In this study, we proposed a support vector machine (SVM)-based method for predicting DHSs, called DHSpred (DNase I Hypersensitive Site predictor in human DNA sequences), which was trained with 174 optimal features. The optimal combination of features was identified from a large set that included nucleotide composition and di- and trinucleotide physicochemical properties, using a random forest algorithm. DHSpred achieved a Matthews correlation coefficient and accuracy of 0.660 and 0.871, respectively, which were 3% higher than those of control SVM predictors trained with non-optimized features, indicating the efficiency of the feature selection method. Furthermore, the performance of DHSpred was superior to that of state-of-the-art predictors. An online prediction server has been developed to assist the scientific community, and is freely available at: http://www.thegleelab.org/DHSpred.html.

## Introduction

Eukaryotic transcription is not only regulated by interactions between transcriptional regulators and *cis*-regulatory DNA elements, but also by chromatin structure, which can affect these interactions. The fundamental unit of chromatins is a nucleosome, which affects transcription; the packing of DNA into nucleosomes inhibits DNA availability to transcriptional regulators [1]. Nucleosome-free regions enhanced with chromatin accessibility, known as DNase I hypersensitive sites (DHSs), have been found predominantly in gene regulatory regions, including promoters, enhancers, and local control regions [2–5]. Since their discovery in 1980, DHSs have been used as markers of regulatory DNA regions [5, 6]. Therefore, mapping of DHSs has become an effective approach for discovering functional DNA elements in noncoding sequences.

DHSs can be identified through southern blotting technique and chromatin immunoprecipitation, followed by microarray hybridization (ChIP-chip) [7, 8]. However, obtaining information on DHSs using the standard southern blot approach is challenging, time-consuming, and error-prone task, and ChIP-chip is expensive and often time-consuming for genome-wide application. Therefore, it is necessary to develop a novel computational method for identifying DHSs. To this end, Noble *et al*. proposed a method based on support vector machine (SVM), using nucleotide composition as a feature vector [9]. Feng *et al.* and Kabir *et al.* also proposed SVM-based predictors, using pseudo-dinucleotide and pseudo-trinucleotide compositions, respectively [10, 11]. All these methods use a similar approach and none of them is available as a web server or stand-alone tool; their practical value is therefore limited, particularly for experimental biologists. Liu *et al.* proposed a predictor called iDHS-EL (identifying DHSs by fusing three different modes of pseudo-nucleotide composition into an ensemble learning framework), which combined three independent random forest (RF)-based predictors, where each one uses different modes of pseudo-nucleotide composition as input features [12]. iDHS-EL is the only publicly available method different from other approaches.

Although these bioinformatics tools showed encouraging results and stimulated research in this area, further studies are needed for the following reasons. (i) The feature space used by the existing methods to construct models is incomplete and not comprehensive. Hence, other potentially useful features remain to be characterized. (*ii*) Biologically significant features are intrinsically heterogeneous and multi-dimensional; however, the existing methods do not employ feature selection techniques to quantify the importance and contribution of features used for the model, leading to only a partial understanding of the sequence-DHS relationships. Due to these deficiencies, other methods are needed to accurately predict DHSs in uncharacterized DNA sequences.

In this study, we developed an SVM-based prediction method for DHSs, called DHSpred (DNase I Hypersensitive Site predictor in human DNA sequences), in which the optimal features are selected using RF (see Figure 1 for an overview of the methodology). The optimal feature candidates are selected using RF from a large set of features, which include k-mer (mononucleotide composition [MNC], dinucleotide composition [DNC], trinucleotide composition [TNC], tetranucleotide composition [TeNC], and pentanucleotide composition [PNC]), dinucleotide physicochemical properties (DPCP), and trinucleotide physicochemical properties (TPCP). In addition to DHSpred, we also developed prediction models using three other machine learning (ML)-based methods (RF, extra-tree classifier [ET], and k-nearest neighbor [k-NN]). Our results showed that the performance of DHSpred was superior to that of three ML-based models and four state-of-the-art predictors.

**Figure 1.**
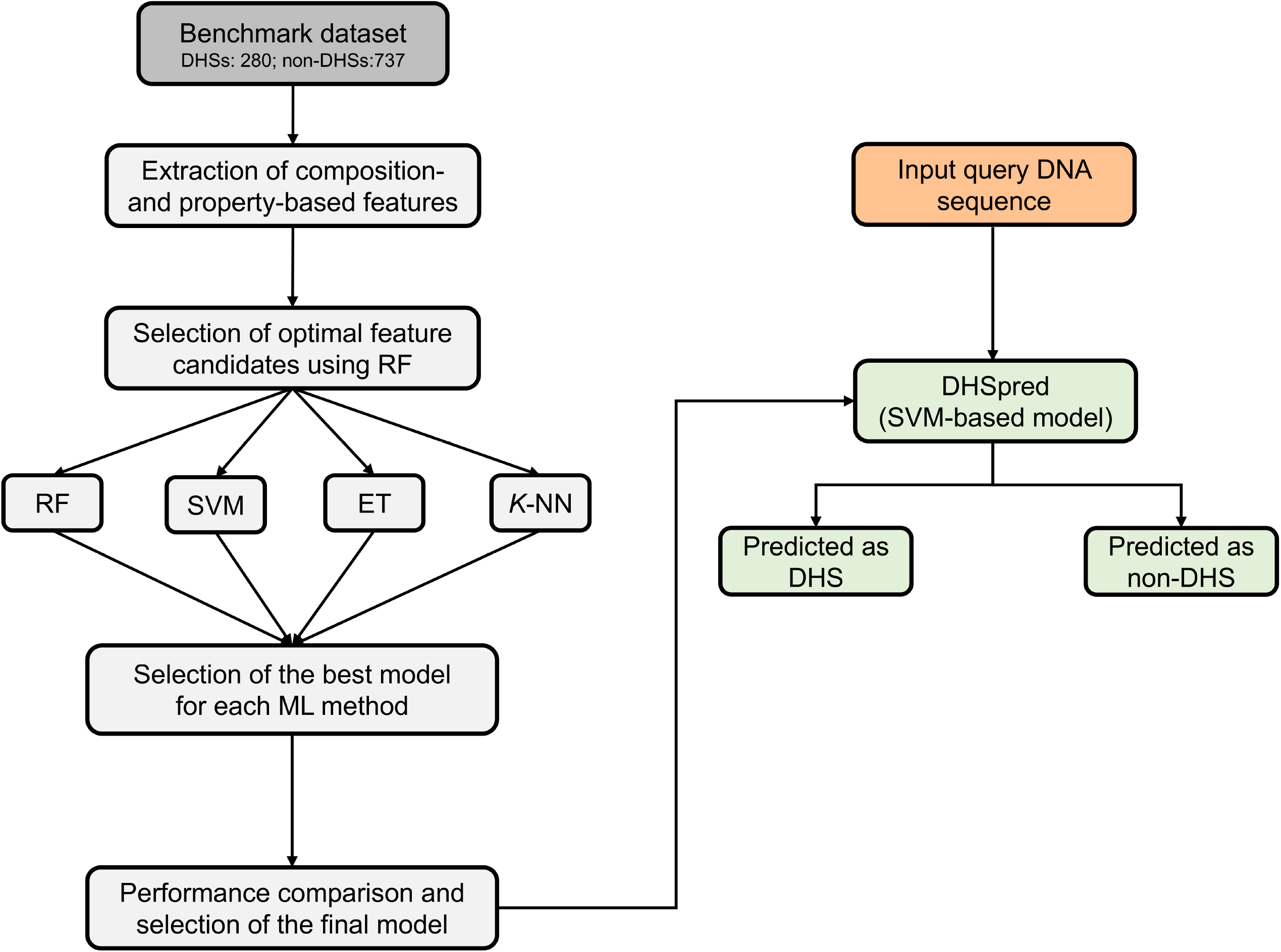
Overall framework of the proposed predictor. Features derived from the DNA sequences, including MNC, DNC, TNC, TeNC, PNC, DPCP, and TPCP, were used as inputs for the RF algorithm to select optimal feature candidates. We then constructed four classifiers based on ML, including RF, SVM, ET, and *k*-NN, using different feature sets, including individual composition, combination of various individual compositions, and feature sets selected from optimal feature candidates. The best model was selected for DHS prediction by comparing the performances of the various models generated in the previous step.

## Results

### Compositional analysis

To understand the human nucleotide bias in DHSs and non-DHSs, we performed compositional analysis of k-mers (MNC, DNC, TNC, TeNC, and PNC) using the benchmarking dataset. MNC analysis revealed that, on average, guanine (G) and cytosine (C) were dominant in DHSs, while adenine (A) and thymine (T) were dominant in non-DHSs (Welch’s *t*-test; *P* ≤ 0.01) (Figure 2A). This trend was also reflected in DNC analysis, where 3 of the top 5 motifs (CG, GC, AT, CC, and TG) abundant in DHSs contained these two (C and G) mononucleotides (Figure 2A), when sorted based on the absolute difference in composition between DHSs and non-DHSs. Furthermore, we observed that 69%, 60%, and 40% of TNC, TeNC, and PNC, respectively, differed significantly between DHSs and non-DHSs (Welch’s ŕ-test; *P* ≤ 0.01). Among them, the top five motifs from TNC (CGC, GCG, CCG, CGG, and GCC), TeNC (CGCC, CCGC, GCGC, GCGG, and GGCG), and PNC (CCGCC, CGCCC, CCGGG, CCCGC, and GGCGG) were from DHSs (Figure 2B, 2C, and 2D) and contained different combinations of C and G. Previous experimental studies have also shown that GC-rich regions showed an open and relaxed chromatin state, which will be convenient for the binding of other macromolecules [13–15]. Results from the compositional analyses suggested that integrating the amino acid preference information would be helpful for differentiating between DHSs and non-DHSs, and so, we used these as input features for ML methods to improve classification. The major advantage of ML methods is their ability to consider multiple features simultaneously, often capturing hidden relationships [16–23].

**Figure 2.**
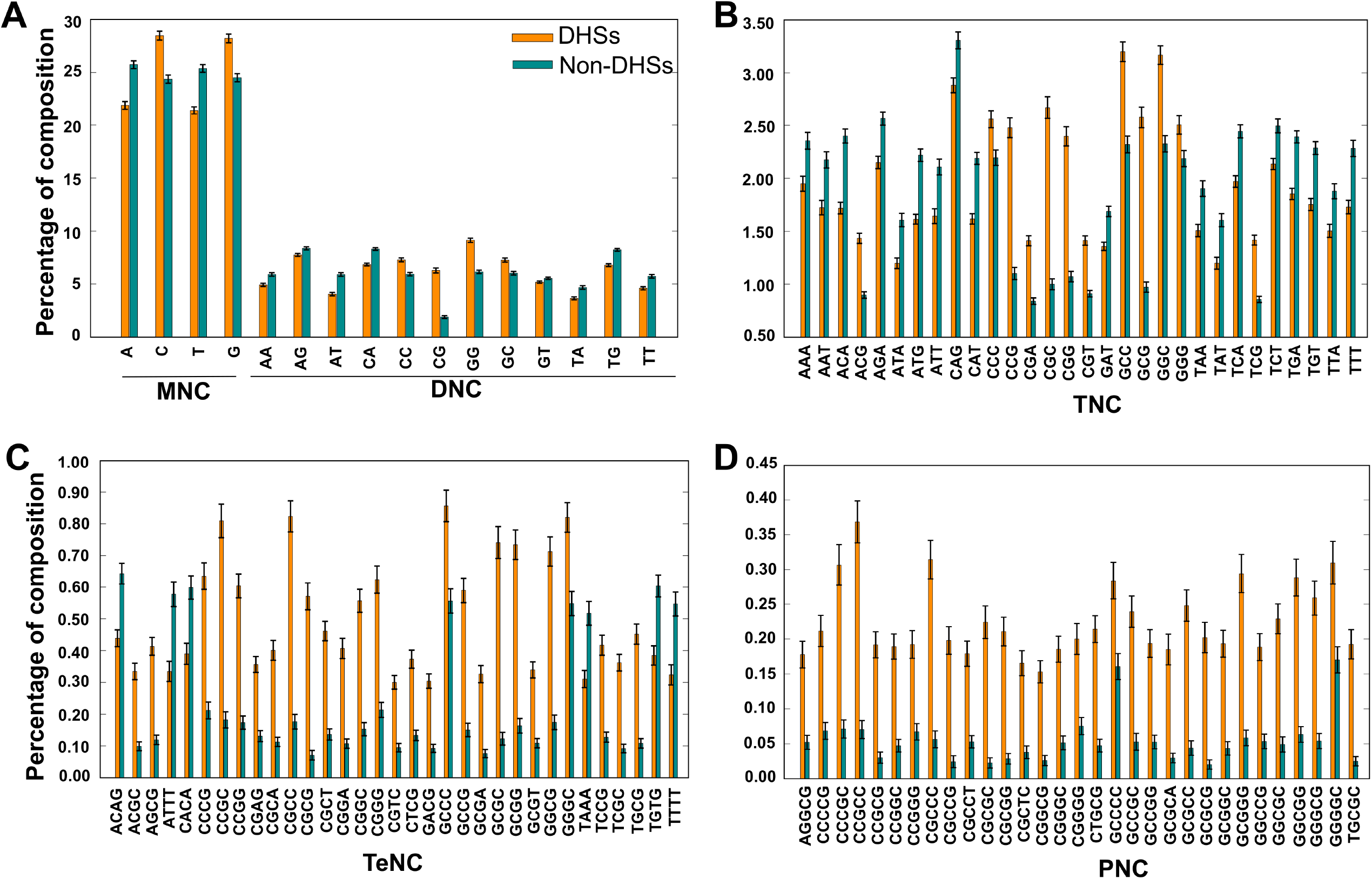
Compositional analysis. (A) MNC and DNC preferences in DHSs and non-DHSs. TNC, TeNC, and PNC preferences between DHSs and non-DHSs is shown in (B), (C), and (D), respectively. In (B), (C), and (D), only compositions with absolute differences between DHSs and non-DHSs of greater than 0.20 are shown.

### Construction of an SVM-based model using the optimal feature set

An SVM-based model was constructed using the optimal feature set, through two steps. In the first step, we employed the RF algorithm and evaluated the importance of 2228 features (MNC: 4; DNC: 16; TNC: 64; TeNC: 256; PNC: 1024; DPCP: 94; and TPCP: 768) for distinguishing DHSs from non-DHSs. Based on a threshold feature importance score (FIS) > 0. 0003, 1139 features were selected as optimal feature candidates (Figure 3A). The percentage of individual contributions to the optimal feature candidates is shown in Figure 3B; most of the contribution was from TPCP, PNC, and TeNC. In the second step, to select more important features, we generated 19 sets of features from the optimal feature candidates using an FIS cut-off (0.0003 ≤ FIS ≤ 0.0021, with a step size of 0.0001). SVM-based prediction models corresponding to these features were then developed. The performance of these 19 prediction models, in terms of Matthews correlation coefficient (MCC), is shown in Figure 3C, where the performance peaked with 174 features (F174). Therefore, we considered this model final, with the optimal feature set. Furthermore, we examined the percentage of individual contributions in the optimal feature set. As shown in Figure 3D, TPCP introduced in this study contributed 51.4%, followed by PNC (20%), TeNC (4%), DPCP (10%), TNC (12.6%), and DNC (2%), indicating that TPCP played a major role in distinguishing DHSs from non-DHSs.

**Figure 3.**
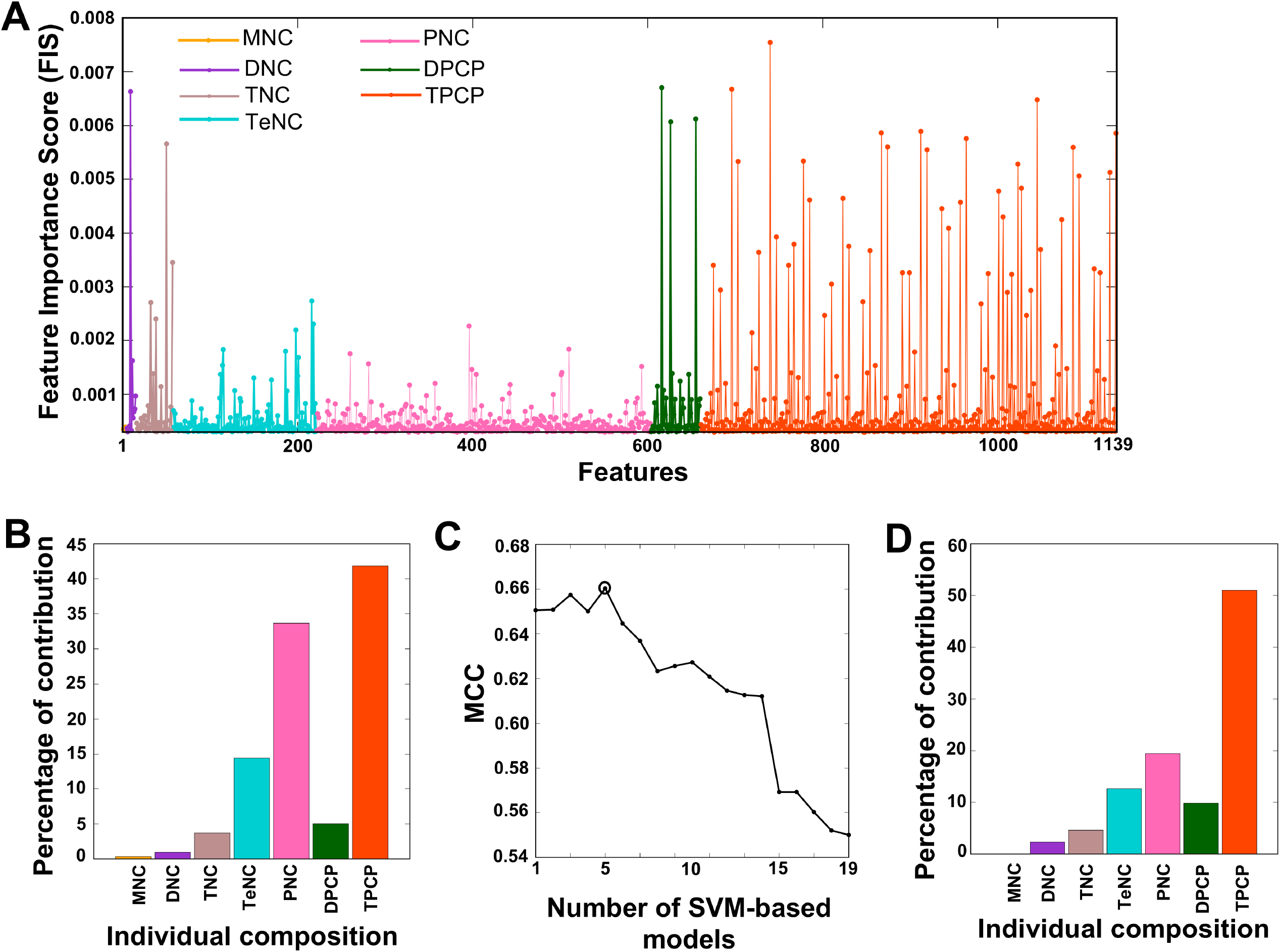
(A) Optimal feature candidates are shown along with their importance scores. The X- and Y-axes represent the features and their importance scores, respectively. The percentage of individual contributions of the optimal feature candidates is shown in (B). (C) The performance of SVM-based models with respect to 19 different sets of features, which were generated from optimal feature candidates using FIS cut-offs. The final selected SVM-based model is circled, and its feature contribution is shown in (D).

Generally, it might be possible for hybrid models (combination of individual compositions) or individual composition-based models to perform better than models developed by rigorous feature selection protocols, such as the one described above. To investigate this possibility, we developed prediction models using individual composition or hybrid models, whose performances in terms of MCC, accuracy, sensitivity, and specificity are shown in Figure 4A. The results showed that the F174-based model was superior to the other models; the F-174 model had an MCC value of 0.660, which was 3% and 2-4% higher than that of the control SVM predictors (non-optimized features or all features) and individual composition-based or hybrid models, respectively. In addition, the number of selected features decreased from 2228 to 174. These results indicated that our feature selection protocol was effective for identifying important and informative features. By removing redundant and less informative features through FIS-based feature selection, we could effectively improve the performance of our model.

**Figure 4.**
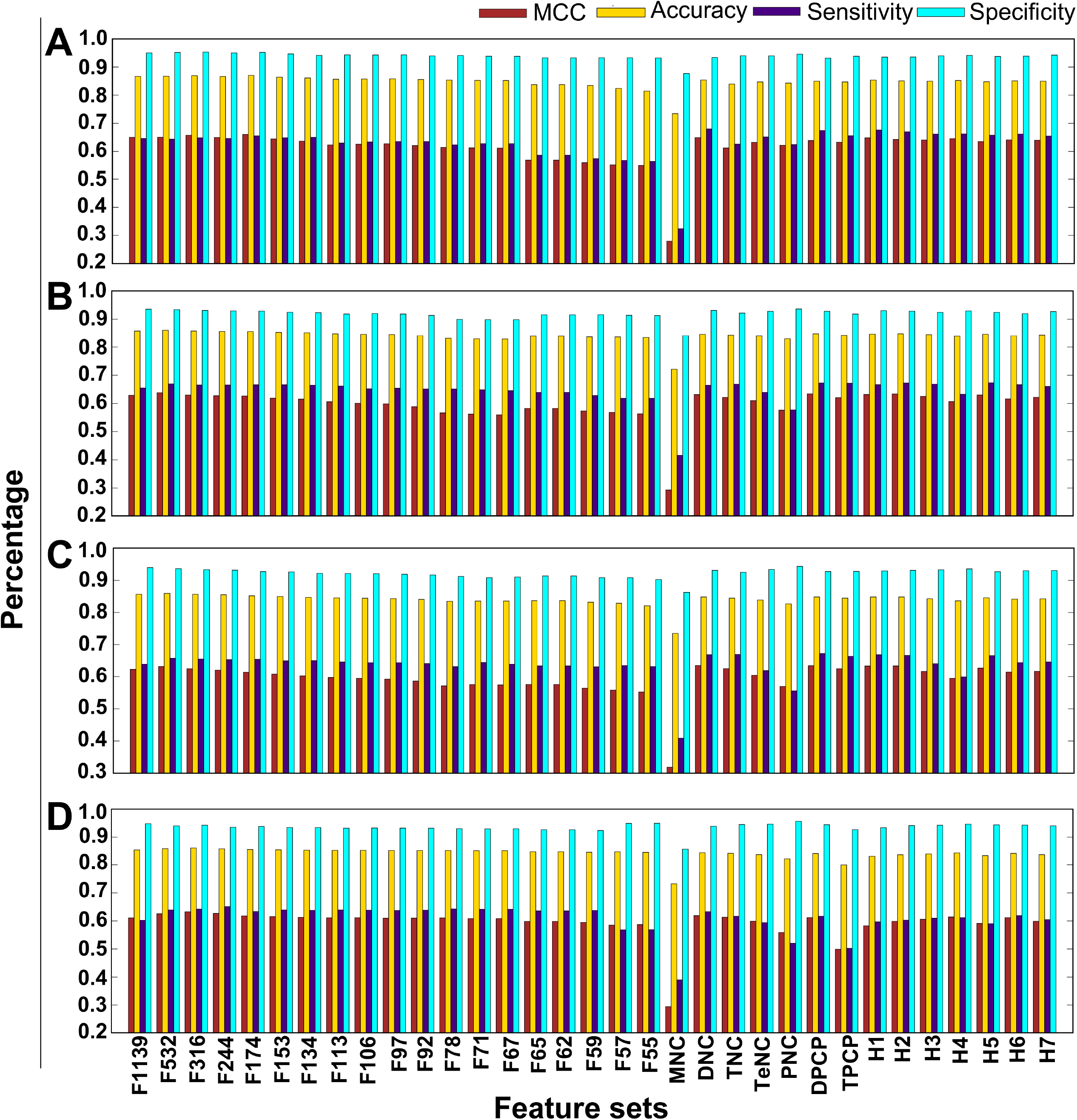
Performance of four different ML-based classifiers. Performance of various classifiers for distinguishing between DHSs and non-DHSs. A total of 33 classifiers were evaluated using five independent 10-fold cross-validation techniques, and their average performances in terms of MCC, accuracy, sensitivity, and specificity are shown. (A) SVM-based performance, (B) RF-based performance, (C) ET-based performance, and (D) *k*-NN-based performance.

### Comparison of three ML-based models with the SVM-based model

In the second step of the previous section, we used three different ML-based methods instead of SVM, including, RF, ET, and k-NN. A detailed description of the development of prediction models using these methods was provided in our recent studies [21, 23]. For each ML-based method, we generated 33 prediction models using different sets of features, including individual composition, hybrid models, and features based on FIS cut-off. The detailed performance of these methods with respect to different feature sets is shown in Figure 4 (B–D). Subsequently, we compared the performance of the three ML-based models with that of the SVM-based model. Interestingly, we observed that the overall performance of the SVM-based model was superior to that of the three ML-based models (ET, RF, and *k-* NN), irrespective of the features used. This result indicated that the SVM method was more suitable for predicting DHSs than other MLs (Figure 5). We then selected the best model from each ML-based method, based on the highest MCC, whose performance is shown in Table 1. Similar to SVM, RF- and k-NN-based methods also showed their best performance when optimal feature sets were selected using different FIS cut-offs. This observation highlights the importance of our feature selection protocol. However, in terms of MCC and accuracy, the SVM-based model was superior to the other ML-based methods by ~2% and 1–2%, respectively.

**Figure 5.**
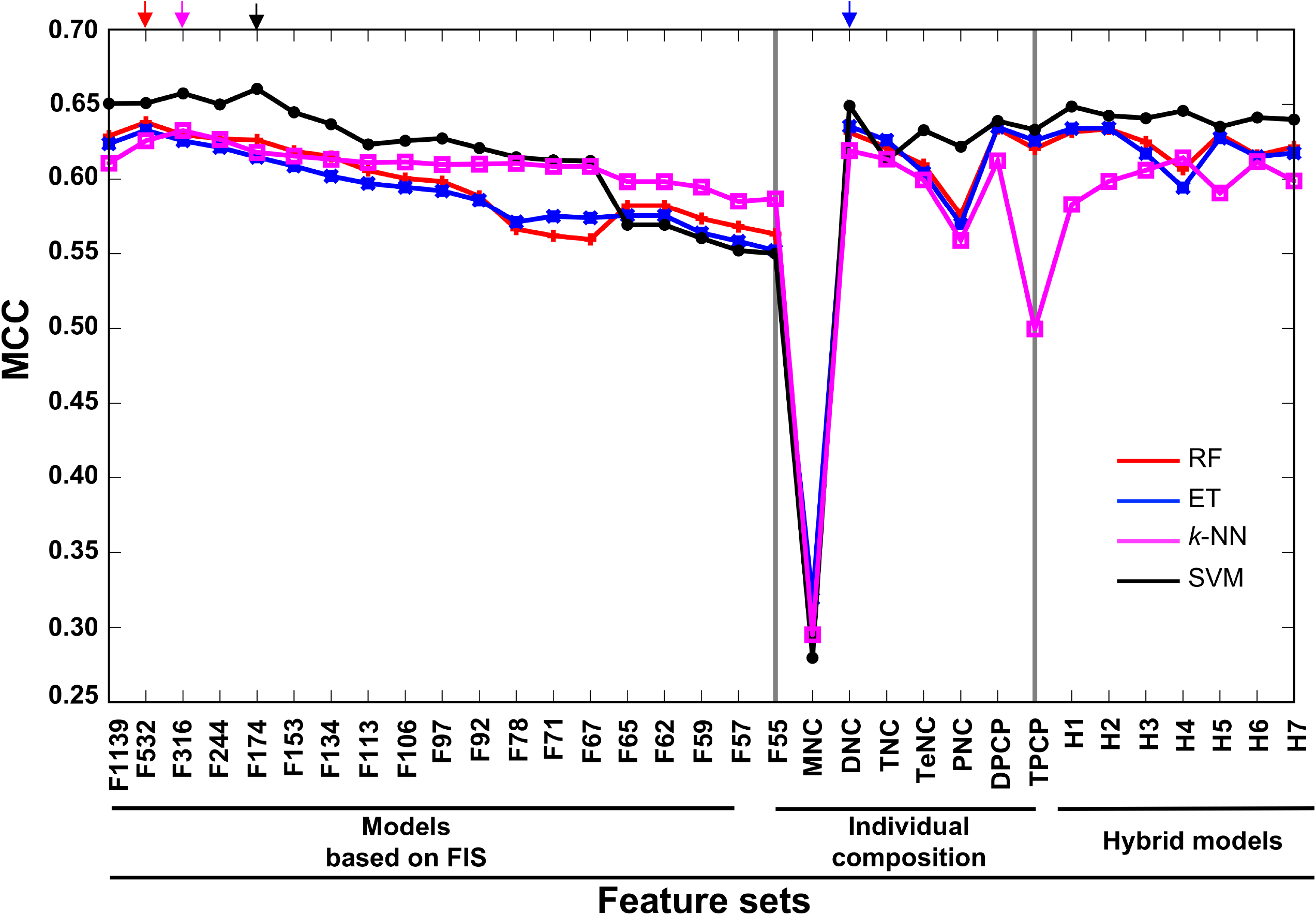
Performance of various prediction models on the benchmarking dataset. X- and Y-axes represent various feature-based prediction models and their performances measured in terms of MCC, respectively. The final selected prediction model for each ML-based method is indicated by arrows in different colors. Each point represents the average of five independent 10-fold cross-validations. The first 19 features are FX, where X is the number of features based on the FIS cut-off. H1: MNC+DNC; H2: H1+TNC; H3: H1+TeNC; H4: H3+PNC+DPCP; H5: DPCP+TPCP; H6: H1+TeNC+PNC+TPCP; and H7: H1+TNC+TeNC+PNC+H5.

**Table 1.** Performance of the best selected model from each ML-based method

| Method | MCC | Accuracy | Sensitivity | Specificity | Pt | Py |
| --- | --- | --- | --- | --- | --- | --- |
| SVM | **0.660** | **0.871** | 0.655 | **0.952** | 0.624 | **0.607** |
| RF | 0.638 | 0.861 | **0.669** | 0.934 | **0.625** | 0.603 |
| ET | 0.635 | 0.848 | 0.668 | 0.931 | 0.622 | 0.599 |
| *k*-NN | 0.632 | 0.860 | 0.642 | 0.943 | 0.605 | 0.585 |

The first column represents the ML-based method developed in this study. The second, the third, the fourth and the fifth, the sixth and the seventh respectively represent the MCC, accuracy, sensitivity, specificity, Pt and Py. Bold font denotes the best result.

### Comparison of state-of-the-art predictors with the SVM-based model

In order to test the quality of the performance of our SVM-based model (DHSpred), it was necessary to compare it to other state-of-the-art methods. Here, we compared it with four such methods (RevcKmer, PseDNC, PseTNC, and iDHS-EL), which were developed from the same benchmark dataset [9–12]. The results are shown in Table 2, in which the methods are ranked according to MCC, which reflects the prediction capability. DHSpred was ranked highest, with MCC, accuracy, sensitivity, specificity, product of sensitivity and specificity (Pt), and product of excess (Py) values of 0.660, 0.871, 0.655, 0.952, 0.624, and 0.607, respectively. For all six metrics (MCC, accuracy, sensitivity, specificity, Pt, and Py), our method had higher scores than the other methods by 2–9%, 1–3%, 0.1–4%, 1–3%, 1.5–6%, and 2–7%, respectively. This result suggested that DHSpred was more effective and robust for identifying DHSs.

**Table 2.** Performance of the proposed DHSpred along with the state-of-the-art methods

| Method | MCC | Accuracy | Sensitivity | Specificity | Pt | Py |
| --- | --- | --- | --- | --- | --- | --- |
| DHSpred | **0.660** | **0.871** | **0.655** | **0.952** | **0.624** | **0.607** |
| iDHS-EL | 0.636 | 0.861 | 0.646 | 0.943 | 0.609 | 0.589 |
| RevcKmer | 0.616 | 0.852 | 0.654 | 0.928 | 0.607 | 0.582 |
| PseTNC | 0.610 | 0.861 | 0.607 | 0.946 | 0.574 | 0.553 |
| PseDNC | 0.571 | 0.837 | 0.611 | 0.923 | 0.563 | 0.533 |

The first column represents the method name employed in this study. The second, the third, the fourth and the fifth, the sixth and the seventh respectively represent the MCC, accuracy, sensitivity, specificity, Pt and Py. Bold font denotes the best result.

The superior efficiency of DHSpred could be attributed to the fact that it selectively incorporates multiple aspects of important information (achieved through systematic feature selection) from k-mer, DPCP, and TPCP, while most other methods use all available information. For instance, iDHS-EL considers all the features from DPCP, instead of merely the important motifs. Although the existing methods could achieve relatively high-level performance, further improvement in performance is limited when handling complicated data samples. Hence, incorporating important features from multiple complementary sources of information (as the DHSpred method does) could allow us to sufficiently capture the differences between true DHSs and non-DHSs, thus improving the predictive performance.

### DHSpred online prediction server

As mentioned in previous publications [20, 24–26], a prediction method along with its web server would be of great practical use for experimentalists [27–33]. To this end, an online prediction server for DHSpred was developed, which is freely accessible at the following link: www.thegleelab.org/DHSpred.html. Users can paste or upload query DNA sequences in the FASTA format. After submitting the input DNA sequences, they can retrieve results in a separate interface. To maximize the convenience for users, step-by-step guidelines are provided in the Supplementary information. To ensure reproducibility of our findings, datasets used in this study can be downloaded from the DHSpred web server.

## Discussion

DHSs are nucleosome-free regions associated with different genomic regulatory elements, including promoters, enhancers, silencers, insulators, and transcription factor binding sites [2–4, 34]. Therefore, identification of DHSs could be an effective approach for discovering functional DNA elements from noncoding sequences. Although many experimental methods have been proposed for identifying DHSs [7, 35], these methods are often expensive, laborious, and time-consuming for genome-wide application. Therefore, it is necessary to develop computational methods for predicting DHSs.

In this study, we developed a novel ML approach, called DHSpred, for predicting DHSs in the human genome. DHSpred combines various informative features from multiple sources, including *k*-mer, DPCP, and TPCP. Although *k*-mer and DPCP features have been used in earlier studies [9, 10, 12], this is the first report in which TPCP features were used. Interestingly, TPCP features contributed to ~52% of the DHS prediction. In ML-based prediction methods, feature selection is an important step, as various data mining tasks have to deal with vast quantities of heterogeneous and possibly redundant features; feature selection has been widely used in various bioinformatics applications to improve the prediction performance [22, 36–38]. Recently, a two-step feature selection protocol was applied in a protein model quality assessment method called SVMQA; the CASP assessors declared this method as the best one for selecting good-quality models from decoys [39]. We applied the same strategy for our current study and developed 132 prediction models using four ML-based methods with different sets of features. Among them, the SVM-based model using the optimal feature set showed the best performance.

In this study, it was demonstrated that DHSpred outperformed state-of-the-art methods (RevcKmer, PseDNC, PseTNC, and iDHS-EL) [9–12] and three other ML-based methods (RF, ET, and *k*-NN) by cross-validations on the same benchmark dataset. The superior performance of our method might be attributed to two important factors: (i) integration of previously reported features and inclusion of novel features that collectively make significant contributions to the performance; (ii) a two-step feature selection protocol to eliminate overlapping and redundant features. Furthermore, our approach is a general one, which is applicable to many other classification problems in structural bioinformatics. It can be readily extended for predicting DHSs in the plant genome.

A user-friendly web interface is also available for researchers to use our prediction method. Indeed, our method is only the second method that is publicly available with high accuracy. Compared to experimental approaches, bioinformatics tools such as DHSpred represent a powerful and cost-effective approach for genome-wide prediction of DHSs. Therefore, DHSpred might be useful for large-scale DHS prediction and facilitate hypothesis-driven experimental design.

## Materials and Methods

According to the Chou’s [40] five-steps guidelines that have been followed in a series of recent publications [12, 20, 24–26, 41–44], to develop a new prediction methods that can be easily used by both experimental scientists and also theoretical scientists, we should follow the following five guidelines: (*i*) construct a valid benchmarking dataset to train and test the prediction model; (*ii*) formulate the biological sequence samples with an effective mathematical expression that can truly reflect their intrinsic correlation with the target to be predicted; (*iii*) introduce or develop a powerful algorithm (or engine) to operate the prediction; (*iv*) properly perform cross-validation tests to objectively evaluate the anticipated accuracy of the predictor; (*v*) establish a user-friendly web-server for the predictor that is accessible to the public. Below, we describe these steps one-by-one.

### Benchmarking dataset

In this study, we utilized the dataset constructed by Noble *et al.* (2005), which was specifically used for studying DHSs [9]. We decided to use this dataset for a number of reasons. (*i*) This dataset is more reliable, as it was constructed rigorously based on experimental data. (*ii*) It is a non-redundant dataset and none of the sequences has high pairwise sequence identity (> 80%) with any other sequence. Hence, this dataset could stringently exclude homology sequences. (*iii*) Most importantly, it will facilitate fair comparison between our method and the existing methods that were developed using the same benchmarking dataset.

Usually, the benchmark dataset comprises a training dataset and a testing dataset. The former is for training a model, while the latter is used for testing one. As pointed out in a comprehensive review [45], there is no need to artificially separate a benchmark dataset into training and testing datasets for validating a prediction method, as long as it is tested by the jackknife or subsampling (K-fold) cross-validation, as the outcome thus obtained is from a combination of different independent dataset tests. Thus, the benchmark dataset taken from Noble *et al.* [9] (2005) can be formulated as

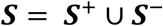

where the positive subset *S*^+^ contained 280 DHS sequences, the negative subset *S*^−^ contained 737 non-DHS samples, and the symbol U denotes union in the set theory. Thus, *S* contained 1017 samples, which can be downloaded from our webserver.

### Feature extraction

The aim of this experiment was to train each ML-based (SVM, RF, ETC, and k-NN) model to accurately map input features extracted from a nucleotide sequence to predict its class *(i.e.,* DHS or non-DHS), which is considered a classification problem. We used nucleotide composition and di- and trinucleotide-physiochemical properties as input features. These features reflect the characteristics of a DNA sequence, from different perspectives, as defined below.

#### k-mer composition

The *k*-mer composition of the nucleotide sequence is calculated using the following equation:

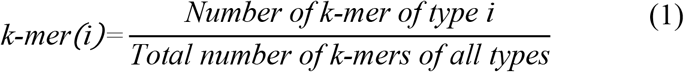

where *i* = 1, 4^*k*^. Therefore, the size of feature vectors for MNC, DNC, TNC, TeNC, and PNC are 4, 16, 64, 256, and 1024, respectively.

#### DPCP

In general, the spatial arrangement of two successive base pairs (bp) can be characterized by six quantities, which include three local translation parameters (shift, slide, and rise) and three local angular parameters (twist, tilt, and roll) [46, 47], which have been successfully used for many sequence-based classifications [12, 49]. In this study, we also utilized DPCP for predicting DHSs. As there are 16 dinucleotides, the total number of features for DPCP would be 6 × 16 = 96.

#### TPCP

Recently, the twelve physiochemical properties (bendability [DNase], bendability [consensus], trinucleotide CG content; nucleosome positioning, consensus [roll], consensus [rigid], DNase I, DNase I [rigid], molecular weight [Daltons], molecular weight [kg], nucleosome, and nucleosome [rigid]) parameters had compiled for DNA trinucleotides [47]. These properties were utilized in the current study for predicting DHSs. As there are 64 trinucleotides, the total number of features for TPCP would be 64 × 12 = 768.

It is to be noted that both DPCP and TPCP parameters were normalized in the range of [0, 1], based on the formula described in our previous studies [50, 51]. To the best of our knowledge, this is the first study in which *k*-mer, DPCP, and TPCP were considered for DHS prediction. Notably, TPCP has never been considered before, although k-mer, DNC, and DPCP are utilized in existing ML-based methods for DHS prediction [9, 10, 12].

### Feature selection

The feature selection protocol is the same as the one used in our recent study [22]. We used a two-step feature selection protocol to identify the most important features for predicting DHSs. In the first step, we applied the RF algorithm to estimate the importance of each feature. A detailed description of how we estimated the importance of input features has been published previously [21, 51, 52]. Briefly, we used all the features as inputs for RF and carried out 10-fold cross-validation on the benchmark dataset. For each round of crossvalidation, we built 10,000 trees, and the number of variables at each node was chosen randomly from between one to 100. The ensemble average of FIS from all the trees is shown in Figure 3.

In the second step, we selected different sets of optimal feature candidates based on an FIS cut-off (0.0001 ≤ FIS ≤ 0.019, with a step size of 0.0001). The sets of features with FIS greater than the cut-off value were selected as the input features for the SVM classifier. For each feature set, we randomly divided our benchmark dataset into ten subsets (~10% of DHSs and non-DHSs in each subset) for each validation step. At each cross-validation step, nine subsets were then merged as the training dataset, in which ML parameters (*C* and γ) were optimized using a grid-search approach to train the model, while the remaining subsets were merged as the test set to validate the built model. This procedure was repeated ten times, and each subset was used in the training and validated in the testing. This 10-fold crossvalidation procedure was repeated five times, resulting in five similar/different ML parameters and performances (see the section on Evaluation metrics). We considered the average performances and median ML parameters as the final stable values, and carried out performance evaluation.

### Support vector machine

SVM is a well-known supervised-ML technique used for developing both classification and regression models based on statistical theory [53]. A detailed description of an SVM has been reported previously [50, 54, 55]. A set of positive (DHS) and negative (non-DHS) samples were represented by the feature vectors *X_i_*(*i*=1, 2,…*N*) with the corresponding labels *Y_i_ ϵ* {0, 1}. To classify the data as DHS or non-DHS, the SVM transforms input samples into one of two classes in a high-dimension feature space and learns an optimal decision boundary or hyperplane using kernel functions. Here, radial basis function (RBF) was set as the kernel function. An RBF-SVM requires the optimization of two critical parameters (C: penalty constant and γ: width). Hence, we optimized these two parameters using a grid search within the following ranges: *C* from 2^−15^ to 2^10^ and *γ* from 2^−10^ to 2^10^ in log_2_ scale. Besides SVM, we also utilized RF and other ML methods (ET and k-NN). These methods were implemented according to the scikit-learn package [56].

### Evaluation metrics

To compare the prediction methods, we evaluated sensitivity (Sn), specificity (Sp), accuracy (ACC), the Matthews correlation coefficient (MCC), product of Sn and Sp (Pt), and property excess (Py) [12, 57]. Among these six metrics, Pt and Py helps in dealing with systems in which the number of negative samples is overwhelmingly greater than that of positive samples, as described by Jin *et al.* [57] (2005) and Yang *et al.* [58] (2005). The conventional formulae for these metrics lack intuitiveness and are not handy for most biologists, particularly MCC. Therefore, Chen *et al.* derived a new set of equations for these metrics [24, 25], based on Chou’s symbols used in a study on protein signal peptide cleavage site [59]. The new formulae for these metrics are given in equation (2).

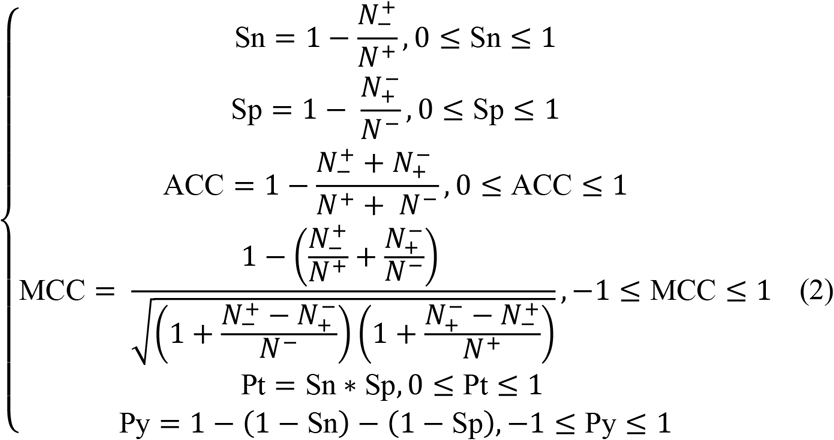

where *N*^+^ is the total number of the DHSs investigated, 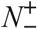 is the number of DHSs incorrectly predicted as non-DHSs, N^−^ is the total number of non-DHSs investigated, and 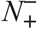 is the number of non-DHSs incorrectly predicted as DHSs. The above set of metrics is valid only for the single-label systems. For the multi-label systems, whose existence has become more frequent in system biology [60] and system medicine [61–63], a completely different set of metrics is needed as defined in [64].

### Development of a prediction server

An online prediction server was developed using hypertext markup language and Java script, with a Python script executing in the backend upon submission of DNA sequences in the FASTA format. Users can submit single or multiple sequences containing standard DNA bp in FASTA format. The DHSpred web server then outputs the results of SVM-based predictions, along with probability values.

## Statistical analysis

The differences in MNC, DNC, TNC, TeNC, and PNC between ORIs and non-ORIs were analyzed using Welch’s *t-test.* The data are presented as mean ± standard error (SE). Statistical differences were considered significant at *p* ≤ 0.01, indicates that the difference is statistically meaningful. All statistical analysis was performed using our own script.

## Abbreviations

DHSpred: DNase I hypersensitive site predictor
DNC: dinucleotide composition
DPCP: dinucleotide physicochemical properties
ET: extra tree classifier
k-NN: k-nearest neighbor
ML: machine learning
MNC: mononucleotide composition
PNC: penta nucleotide composition
RF: random forest
SVM: support vector machine (SVM)
TeNC: tetra nucleotide composition
TNC: trinucleotide composition
TPCP: trinucleotide physicochemical properties

## Author contributions

Conceived and designed the experiments: BM, GL. Performed the experiments: BM. Analyzed the data: BM, THS, GL. Contirbuted reagents/mateials/software tools: THS, GL. Wrote paper: BM, GL.

## Acknowledgments and funding

This work was supported by the Basic Science Research Program through the National Research Foundation (NRF) of Korea funded by the Ministry of Education, Science and Technology (2015R1D1A1A09060192), Priority Research Centers Program through the National Research Foundation of Korea (NRF) funded by the Ministry of Education, Science and Technology (2009-0093826), and the Brain Research Program through the National Research Foundation of Korea (NRF) funded by the Ministry of Science, ICT & Future Planning (2016M3C7A1904392).

